# No facilitatory effects of transcranial random noise stimulation on motion processing: A registered report

**DOI:** 10.1101/2025.03.18.643903

**Authors:** Grace Edwards, Ryan Ruhde, Mica B. Carroll, Chris I. Baker

**Author notes:** Equal contribution (Grace Edwards, Ryan Ruhde, Mica B. Carroll). **Corresponding author** Grace Edwards, New address: The National Institutes of Health, NIMH, Laboratory of Brain and Cognition, 10 Center Drive, Room 4C104, Bethesda MD 20892-1366.

## Abstract

Non-invasive brain stimulation (NIBS) techniques have the potential to demonstrate the causal impact of targeted brain regions on specific behaviors, and to regulate or facilitate behavior in clinical applications. Transcranial random noise stimulation (tRNS) is one form of transcranial electric stimulation (tES) in which an alternating current is passed between electrodes at random frequencies. High-frequency tRNS (hf-tRNS) is thought to enhance excitability and has been reported to have facilitatory effects on behavior in healthy and clinical populations. Due to the potential application of tRNS, clear demonstrations of the efficacy and replicability of stimulation are critical. Here, we focused on replicating the facilitatory effect of hf-tRNS over the human middle temporal complex (hMT+) on contralateral motion processing, initially demonstrated by Ghin et al. (2018). In this prior study, the improvement in performance was specific to global motion processing in the visual field contralateral to stimulation. The motivation to replicate this effect was reinforced by the well-supported hypothesis that hMT+ is critical for contralateral global motion processing. However, our results indicated that hf-tRNS does not improve motion discrimination. Specifically, we were unable to replicate a contralateral global motion processing facilitation following hf-tRNS to hMT+. In our within-subject controls, we also found no difference between hf-tRNS to hMT+ on contralateral global motion processing in comparison to sham stimulation, or in comparison to hf-tRNS to the forehead. While it remains possible that our lack of replication could be due to minor changes in the protocol from the original Ghin et al., study, for hf-tRNS to become a widely applied method, the modulatory effect of hf-tRNS should be robust to slight adjustments to the procedure.

## Introduction

Targeting a specific brain region with non-invasive brain stimulation (NIBS) clarifies that region’s functional and causal role in subsequent behavioral change. By regulating behavior, based on application and type, NIBS has potential both as an experimental research method and in clinical applications. Transcranial random noise stimulation (tRNS) is a variety of transcranial electric stimulation (tES) in which an alternating current is passed between electrodes at random frequencies (Terney et al., 2008). tES has been shown to be more cost-effective, accessible, and well-tolerated than other NIBS, such as transcranial magnetic stimulation (TMS; Westwood, 2020). tRNS is a less commonly used method of tES in comparison to transcranial direct current stimulation (tDCS) and transcranial alternating current stimulation (tACS), however the application of tRNS is increasing rapidly. tRNS, unlike tDCS, has polarity independence and no uniform electrical field direction, meaning that it can target multiple brain regions simultaneously (van der Groen et al., 2022). High-frequency tRNS (hf-tRNS) is thought to enhance excitability and predominantly causes facilitatory effects on behavior, in some cases culminating in a performance increase of ∼30% (Contò et al., 2021). tRNS has been demonstrated to enhance global motion direction discrimination (Ghin et al., 2018), fluid intelligence (Brem et al., 2018), numerosity (Cappelletti et al., 2013), arithmetic ability (Snowball et al., 2013), and spatial attention (Contò et al., 2021).

tRNS provides the research community with a method to demonstrate the causal impact of targeted brain regions on specific behaviors, with applications in the clinical community becoming increasingly evident. For example, Herpich et al. (2019) found that tRNS over the visual cortex improved visual motion perception in both healthy controls and cortically blind patients over the course of ten days. Further, they found that the effect persisted for at least six months without further stimulation, suggesting that long-term plastic change in sensory processing was responsible for the observed effect.

Along with evidence of successful neuromodulation, however, there are examples of a failure to replicate these effects. For example, Romanska et al. (2015) found that tRNS over lateral occipitotemporal cortices improved performance on a facial identity perception task. However, Willis et al. (2019) failed to replicate this effect. Changes in methodology, such as the use of a different face stimulus set or the application of a different stimulation intensity (1.5 mA as opposed to 1 mA) may have contributed to the lack of replicability. However, the variability in outcome highlights the critical importance of replication studies in highlighting the flexibility of the procedure, especially given the potential applications of tRNS. For tRNS to be a widely applied method, the modulatory effect of tRNS should generalize despite slight alterations to the procedure.

Here, we attempted to replicate the facilitatory effect of hf-tRNS over the human middle temporal complex (hMT+) on contralateral motion processing, initially demonstrated by Ghin et al. (2018). Specifically, they reported that hf-tRNS over the hMT+ and vertex (Cz) enhanced sensitivity to global motion in a dot array stimulus. The improvement in performance was specific to global motion processing in the visual field contralateral to stimulation. No such stimulation impact on contralateral global motion processing was found for anodal- or cathodal-tDCS, or sham stimulation.

The motivation to replicate the hf-tRNS effect in Ghin et al. (2018) was reinforced by the well-supported hypothesis that hMT+ is specific to contralateral global motion processing (Strong et al., 2017; Ajina et al., 2015; Braddick et al., 2001). Many NIBS studies have investigated the effects of stimulation over hMT+ on motion processing and have found reliable effects on neural activity and task performance (e.g. Antal et al., 2004; Antal et al., 2012; Campana et al., 2016; Pavan et al., 2019). Combined, these studies demonstrate that stimulation over hMT+ has consistent effects. For example, Antal et al. (2004) tested the impact of tDCS (anodal and cathodal) on hMT+, V1, and the motor cortex during a visuomotor coordination task and a motion direction discrimination task. The authors found that only anodal-tDCS over hMT+ improved performance on these tasks. TMS over hMT+ has also been shown to modulate motion processing. For example, Laycock et al. (2007) found that single pulse TMS to hMT+ 158 ms after stimulus onset of a motion direction discrimination task disrupted task performance. Similarly, McKeefry et al. (2008) found that repetitive TMS over hMT+ and V3A caused deficits in speed discrimination, while TMS over adjacent areas and V1 did not. Finally, Campana, Maniglia, and Pavan (2013) found that repetitive TMS over hMT+ reduced the duration of dynamic and static motion after-effect.

Motion processing is predominantly lateralized, which enables the comparison of stimulation on the left and right visual fields as a within session control. A tDCS study found that cathodal-tDCS over left hMT+/V5 reduced a noisy signal while anodal-tDCS boosted a weak signal in the contralateral visual field only during a motion coherence task (Battaglini, Noventa, and Casco, 2017). Furthermore, repetitive TMS over left hMT+ impaired performance in a multiple object tracking task in the contralateral visual field only (Chakraborty et al., 2021). Due to strong theoretical support and evidence across NIBS techniques for the highly reliable effects of stimulation over hMT+, our study provided an ideal test for the replicability of hf-tRNS effects.

We attempted to replicate and extend Ghin et al.’s (2018) findings by combining Experiments 1 and 2 in their paper. In Experiment 1, the authors employed a within-subjects design with three types of tES (cathodal-tDCS, anodal-tDCS, hf-tRNS) and sham stimulation. Each participant (n=16) underwent each type of stimulation over the course of four, non-consecutive sessions in which the active electrode was placed over left hMT+ and the reference electrode was placed over Cz. The authors applied stimulation during a motion direction task (i.e. online stimulation) which required participants to determine the overall direction of random dot kinematograms (RDK’s), or fields of moving dots, during an eight alternative forced choice task. RDK’s were either shown to the left or right of a fixation dot in the middle of the screen. An adaptive maximum likelihood procedure (MLP) was used to determine the coherence threshold of the stimuli in each trial at which the participants performed at 70% accuracy within 18 minutes (Grassi and Soranzo 2009). The authors found that only the hf-tRNS condition lowered the coherence threshold in the right visual field, contralateral to the active electrode placed over the left hMT+ (ipsilateral > contralateral = 10.51%). Anodal- and cathodal-tDCS had no impact on performance.

To verify that the effects of Experiment 1 were location-specific, Ghin et al. (2018) stimulated two control sites in Experiment 2. In the first control (n=12), the active electrodes were placed over Cz and the left forehead. This control examined if stimulation over Cz alone affected motion direction discrimination in Experiment 1. In the second control (n=12), the active electrodes were placed over Cz and the left primary visual cortex (V1) to determine if visual cortex stimulation was sufficient to improve motion direction discrimination. No significant effects on performance in the motion direction discrimination task were found in either control (forehead: ipsilateral > contralateral = 1.17%; V1: ipsilateral > contralateral = 1.58%), apparently confirming the hypothesis that the observed effects of stimulation were specific to hMT+. However, the authors did not directly compare the hMT+ condition with the V1 or forehead control conditions.

Although we aimed to replicate the contralateral impact of hf-tRNS over left hMT+ from Ghin et al., (2018), we did not register an exact replication. We proposed to include only a selection of the original conditions, and introduced extra within-subject stimulation controls in our design. We combined Experiments 1 and 2 from Ghin et al. (2018) to create a three condition within-subjects design: from Experiment 1, we retained the hf-tRNS condition over left hMT+ to Cz, and sham condition over left hMT+ to Cz. Due to the lack of stimulation impact following anodal- or cathodal-tDCS, we did not register to replicate these effects. In order to perform the direct comparison of hMT+ with another active hf-tRNS control site, from Experiment 2, we adopted the hf-tRNS over the left forehead to Cz stimulation control rather than left V1 stimulation. Simulations of hMT+, forehead, and V1 stimulation (all in combination with Cz as the second electrode), demonstrated an overlap of stimulated posterior areas in the hMT+ and V1 simulations (see Appendix A). Due to the overlap of stimulated areas, we did not use hf-tRNS over V1 to Cz as an additional active control. Although we maintained the original predetermined coordinate stimulation procedure of Ghin et al. (2018), we also localized each participants’ hMT+ using lateralized radial moving dots (Strong et al., 2017). We used the localization to calculate the overlap between the predetermined coordinate stimulation and individual neural activity in response to lateralized motion processing. We predicted the measure of overlap between e-field flow and neural activity would correlate with stimulation effect on behavior. By maintaining the stimulation approach of Ghin et al. (2018), we ensured our replication was closer to the original study. All methodological differences are outlined in Table 1 at the end of the methods section.

**Table 1:**
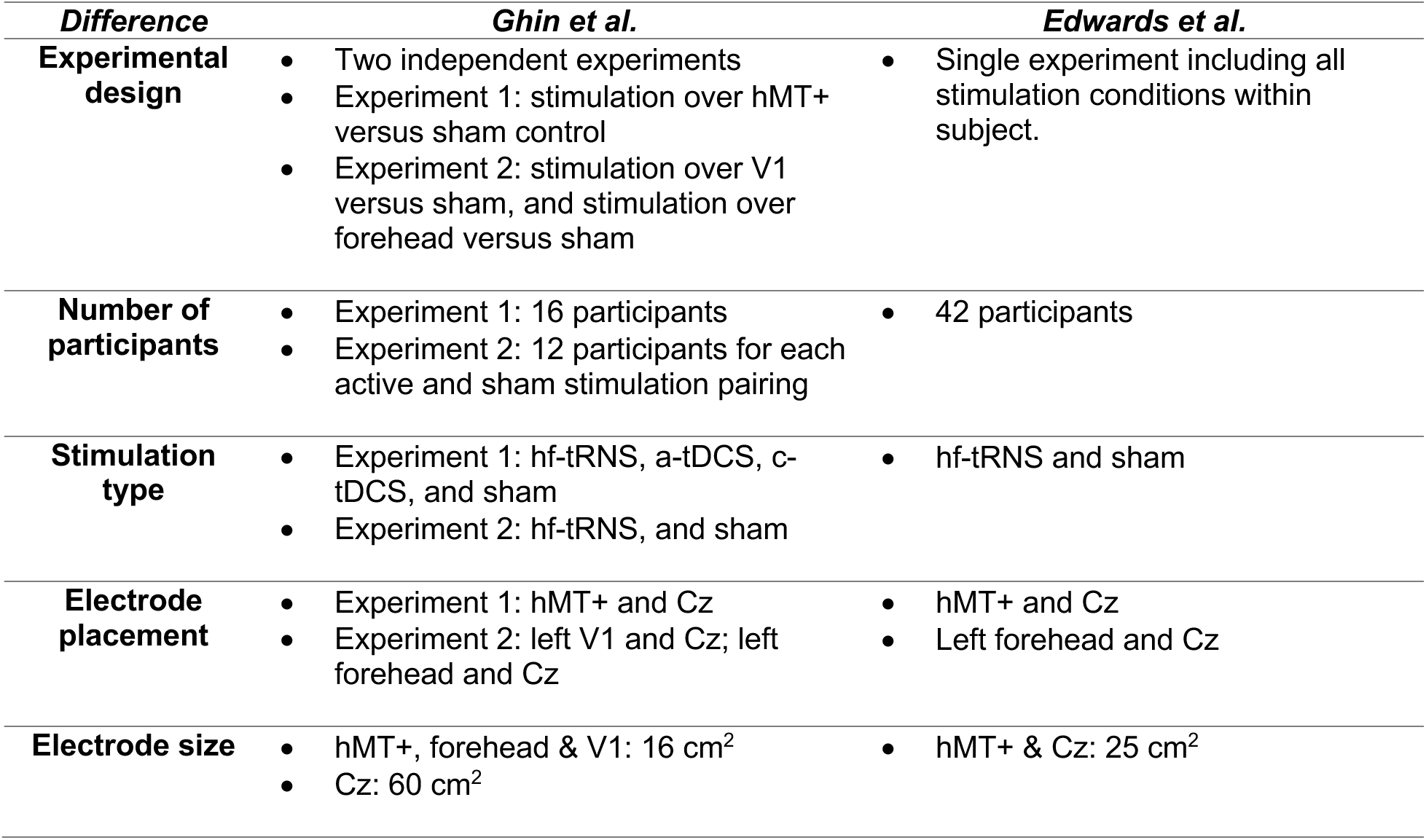
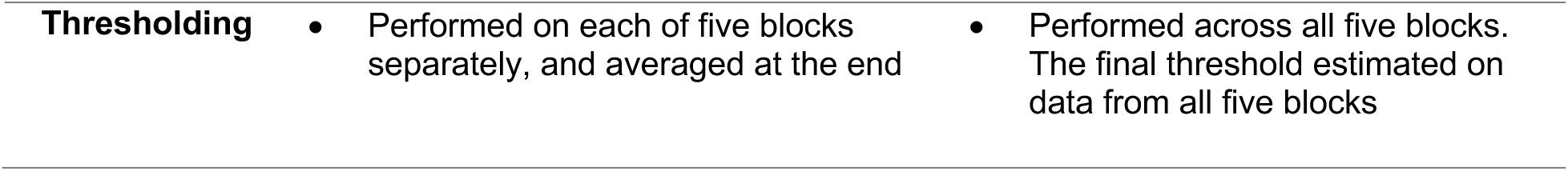
The methodological differences between the original Ghin et al., 2018 study and our proposed replication. Particular attention paid to Experiments 1 and 2 of the Ghin et al., 2018 study which we propose to replicate.

**Table 2:**
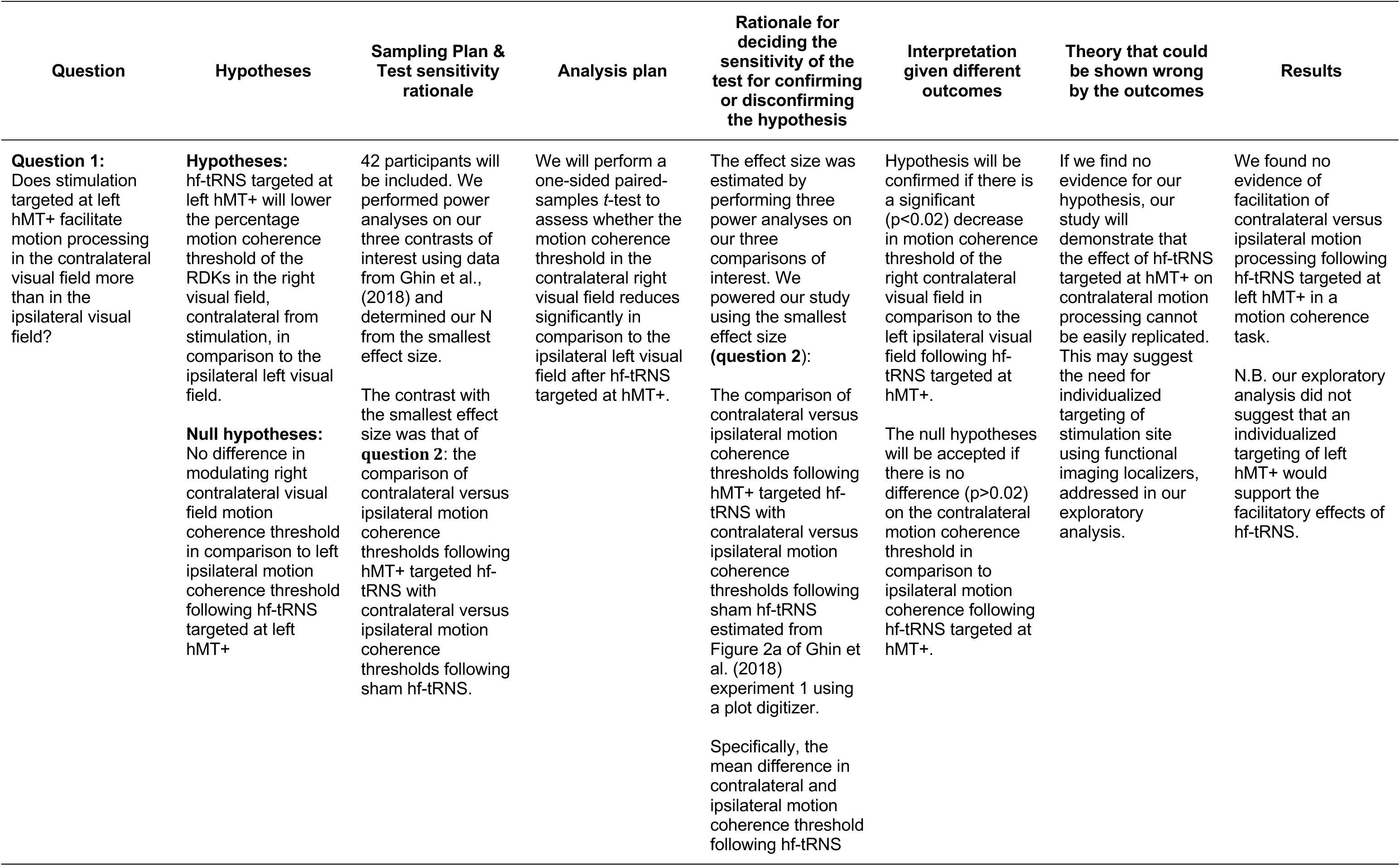
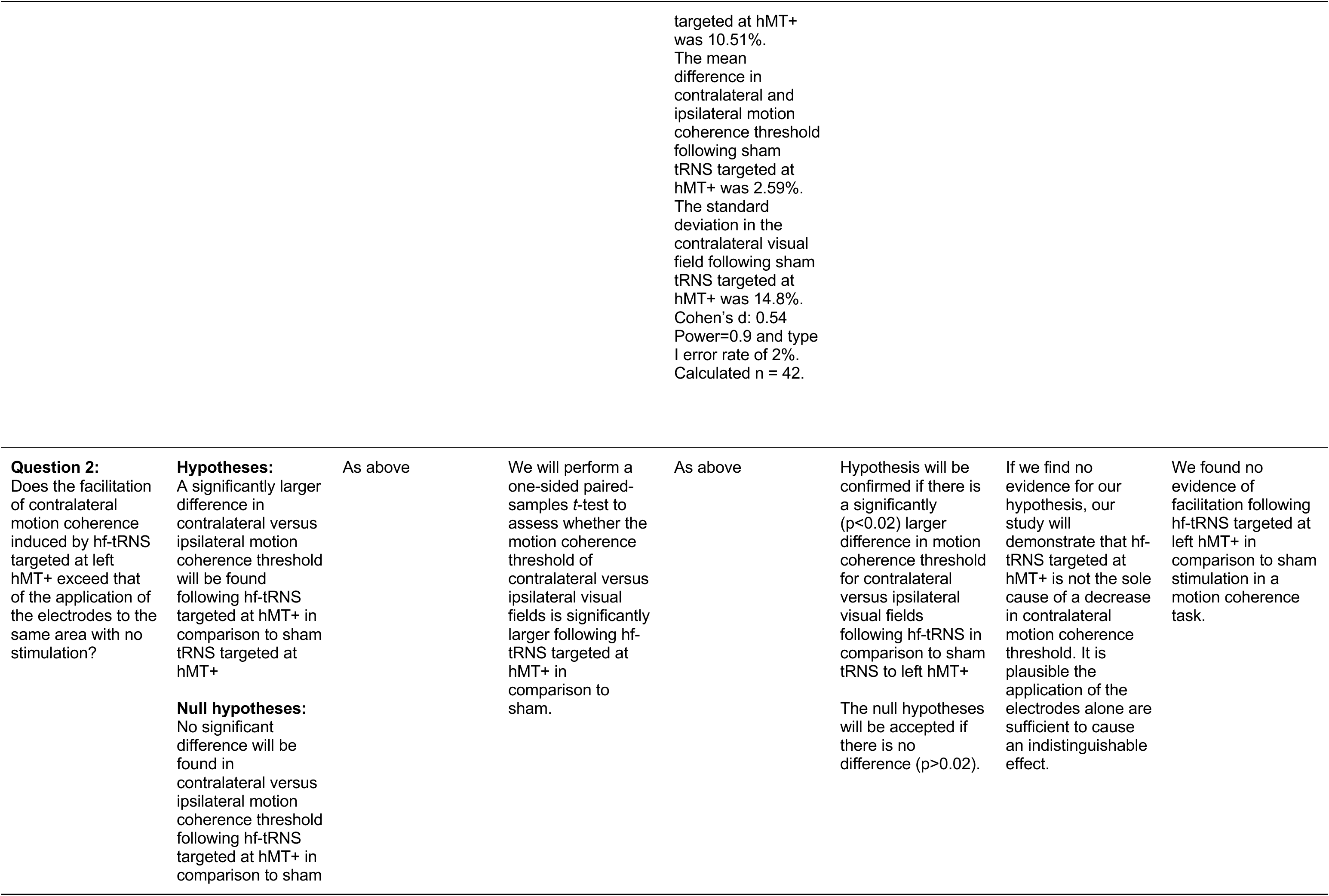
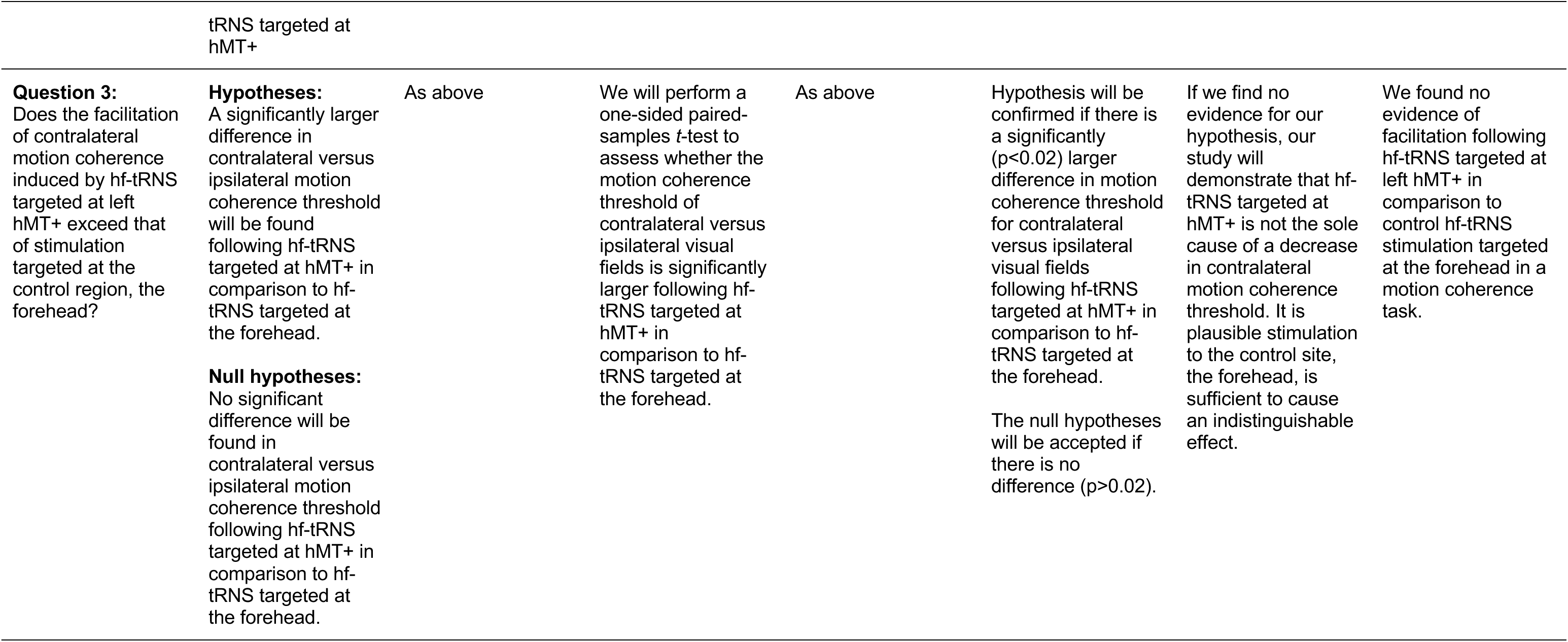
Study Design Table. To conclude contralateral facilitation of motion coherence processing is caused specifically by hf-tRNS targeted at hMT+, all three hypotheses need to be supported.

We had three main hypotheses: 1) We hypothesized that we would replicate the facilitatory effect of hf-tRNS to hMT+ on contralateral global motion processing in comparison to ipsilateral global motion processing. 2) We expected the facilitation for contralateral in comparison to ipsilateral motion processing would be larger for hMT+ in comparison to sham. 3) We expected the facilitation for contralateral in comparison to ipsilateral motion processing would be larger for hMT+ in comparison to forehead stimulation.

We also performed an exploratory analysis to examine if an increased overlap between the predetermined hMT+ targeted stimulation coordinates and neural response to processing lateralized motion predicted an increased impact of hMT+ targeted stimulation. The Stage 1 registered report received in-principle acceptance and is available on OSF (https://osf.io/bce7u).

## Methods

### 1. Participants

After exclusions, forty-two participants took part in the study, determined by the largest number of participants needed for the three effect sizes of interest using the data from Ghin et al. (2018). We performed three power analyses on the three comparisons of interest (See Appendix B). We acknowledge that using a central estimate of effect size from a standard published (not registered report) study may risk an overestimation of effect size. The power analyses for which we estimated our sample size used a between-subjects standard deviation from Ghin et al. (2018), providing a conservative estimate of variance for our within-subjects comparisons. Based on these power analyses we predicted 42 participants to be the number of participants necessary to achieve a 0.9 power to test our minimum effect size of interest.

Participants were only excluded from analysis if there was an equipment failure, or if the participant failed to attend all sessions. We continued to recruit participants until we had 42 valid datasets after exclusion. Four participants were excluded due to equipment failure. Eleven participants were excluded due to failure to attend all sessions. Participants were not excluded due to outliers as the counterbalancing of the three stimulation sessions could have led to spurious outliers as a result of learning. The participants were screened for normal or corrected-to-normal visual acuity and the ability to meet a 70% threshold during the constant thresholding procedure (see Appendix C). This study was approved by the National Institutes of Health Institutional Review Board. We obtained written informed consent from each participant before taking part in the study, for which they received monetary compensation.

### 2. Apparatus

We displayed the stimuli on a ViewPixx monitor (21.25 x 12 inches) with a refresh rate of 60 Hz. Screen resolution was 1920 x 1080 pixels, with participants 25.65 inches from the screen. The minimum luminance of the screen was 0.51 cd/m^2^, the maximum 84.35 cd/m^2^, and the mean 42.58 cd/m^2^. We generated stimuli using Matlab R2016b with Psychtoolbox-3.

### 3. Stimuli

#### A. Random Dot Kinematograms (RDKs)

We used random dot kinematograms (RDKs) presented in either the left or right visual field, the same stimuli as in Ghin et al. (2018) (Figure 1). Each RDK consisted of 150 white dots (0.12 deg diameter) and was presented in a circular aperture (8 deg diameter; 3 dots/deg^2^ density). The center of the aperture was placed 12° to the left or right of fixation. Dots drifted at a speed of 13.3 deg/s, and either lasted for 47 ms or reached the edge of the aperture before they vanished and were replaced by new dots at a randomly selected position within the circular aperture. This maintained dot density. Dots appeared asynchronously and belonged to one of two categories: signal dots and noise dots. Signal dots only moved along one of the eight cardinal trajectories, while noise dots appeared at randomly selected locations within the aperture on each new frame. Further, dots had an equal probability of being selected as a signal dot or noise dot to minimize motion streaks. Each RDK lasted for 106 ms; the short duration of the stimuli was designed to prevent attentional tracking of motion direction and eye movements towards the RDK’s. At the end of each RDK presentation, the participants had three seconds to determine the overall direction of the moving dots in an eight-alternative forced choice task.

**Figure 1.**
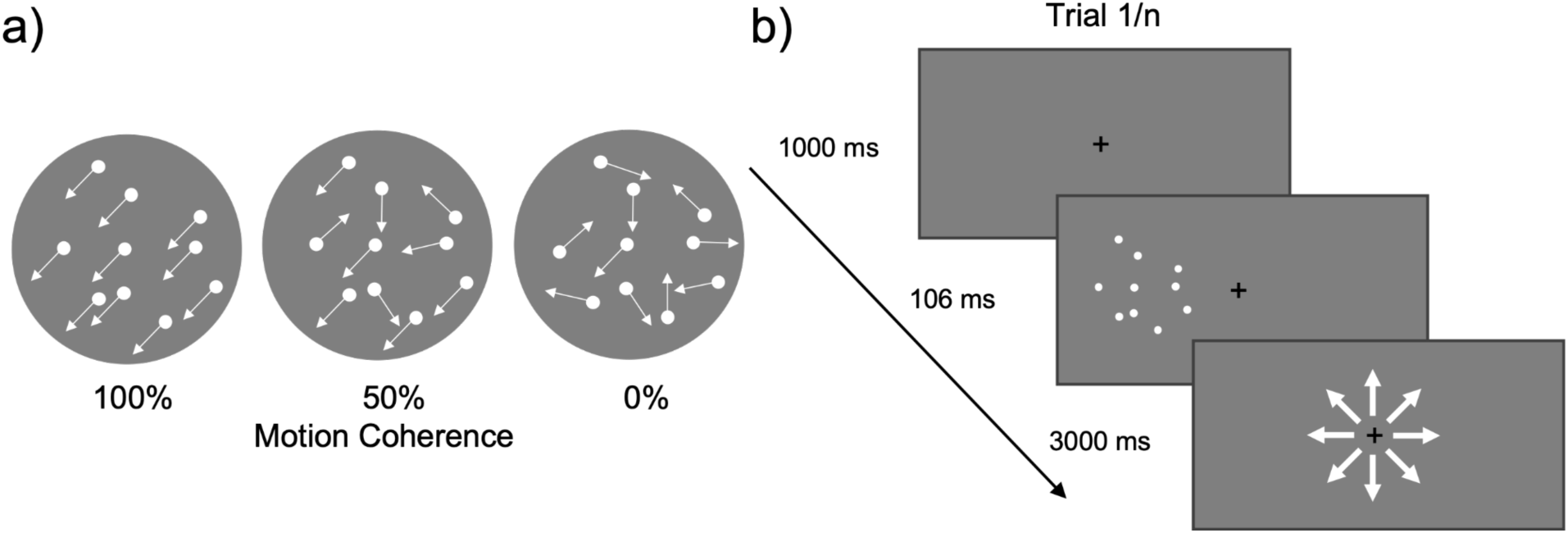
Illustration of RDK’s. a) RDKs with motion coherence of 100%, 50%, and 0%. B) A depiction of one trial of the motion discrimination task. RDKs were randomly presented 12 dva to the left or right of fixation on each trial.

#### B. Thresholding

In each visual field we estimated the motion coherence threshold corresponding to 70% correct motion discrimination. Motion coherence was reported as the percentage of dots moving in the same direction that enabled successful motion discrimination. The motion coherence was determined in each visual field using an interleaved adaptive maximum likelihood procedure (MLP; Grassi and Soranzo, 2009). MLP is a parametric adaptive thresholding procedure, which is less time consuming that non-parametric procedures. Adaptive means the percentage motion coherence for trial n+1 was selected based on the participant’s response to the previous n trials. Adaptive procedures therefore maximized the number of trials presented close to threshold.

The threshold was estimated in five steps:

1. Collection of the n-th trial response
2. Fit the previous responses of n trials to hypothesized psychometric functions.
3. Selection of the psychometric function maximizing the likelihood of the previous n trials.
4. Using the selected psychometric function, estimate the percentage motion coherence to present 70% performance accuracy.
5. Present the motion coherence percentage for the subsequent trial. The final estimated motion coherence percentage was the final coherence threshold.

Our psychometric functions

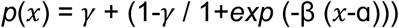

had a fixed slope (β) of 1/2, a fixed baseline (γ) of 12.5% in accordance with chance level in an eight-alternative forced choice paradigm, and a target threshold (*p*) of 70%. The psychometric functions only differ by midpoint (ɑ), which range across 150 values enabling presentation of 0-100% motion coherence at 70% accuracy. On each trial the MLP determined which of the hypothesized 150 psychometric functions best fit the data from the previous trials. The MLP code was adapted from Grassi and Soranzo (2009).

The coherence threshold was estimated using MLPs across all 160 trials, presented over five blocks, during stimulation. Ghin et al. (2018) ran a MLP for each block and averaged across all five. In order to provide the MLPs with maximum trials, we thresholded across all five blocks combined (methodological differences listed in Table 1).

#### C. Localizer stimuli

The left hMT+ was identified for each participant using a fMRI localizer similar to the one described in Strong et al. (2017) (see Figure 2). Across two localizer runs, participants viewed 32 blocks with apertures of moving or static dots presented 12° to either the left or right of fixation. Every four blocks, the participants had a fixation block. Each block was presented for 15 s. In the moving and static blocks, the 10° diameter aperture contained 300 white dots (∼0.2° diameter) presented on a black background. In the movement blocks, the dots moved at 7°/s radially alternating inwards and outwards. The participants performed a task at fixation, pressing a button every time the fixation point turned pink.

**Figure 2.**
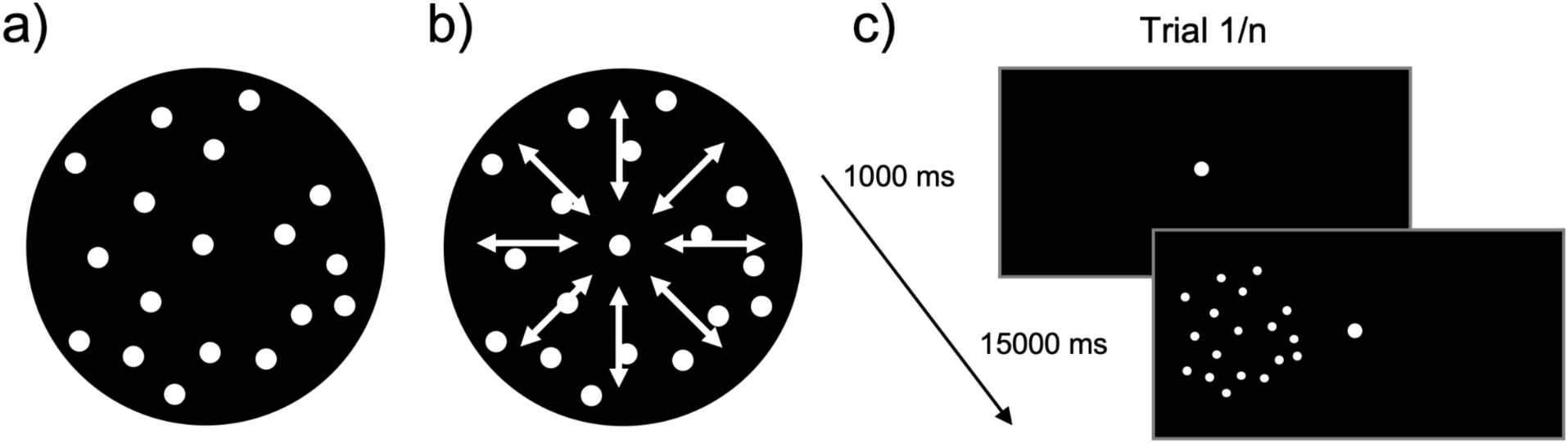
Illustration of localizer stimuli. A) represents a static dot stimulus, b) represents a radial dot motion stimulus. C) A depiction of one block of the localizer. If F = fixation, MR = motion right of fixation, ML = motion left of fixation, SR = static right of fixation, SL = static left of fixation, the run presentation order was as follows: Run 1: F, MR, SR, SL, ML, F, SR, ML, MR, SL, F, SL, MR, ML, SR, F, ML, SL, SR, MR, F. Run 2: F, ML, SL, SR, MR, F, SL, MR, ML, SR, F, SR, ML, MR, SL, F, MR, SR, SL, ML, F.

### 4. fMRI acquisition

We used a 3.0T GE 750 MRI scanner in the fMRI facility in the Clinical Research Center on the National Institutes of Health campus in Bethesda, MD to collect the hMT+ localizer and MPRAGE anatomical scans. Gradient echo pulse sequences were used with a 32-channel coil to measure blood oxygenation level-dependent (BOLD) signal during the functional localizer (TR = 2000 ms, TE = 39, Frequency FOV = 22.5 cm, 50 slices, 2.5 x 2.5 x 2.5 mm^3^, interleaved slice acquisition). A high-resolution T1-weighted anatomical was also collected (208 slices, TR=7, TE= Min full, Frequency FOV = 25.6 cm, flip angle = 8°).

### 5. fMRI analysis and localization of the hMT+

The functional and anatomical MRI data was preprocessed and analyzed using AFNI (Cox 1996). During preprocessing of the functional images, we motion-corrected the data after removing the dummy scans from the analysis (n=3), allowing stabilization of the magnetic field. We then aligned our functional data to each individual’s anatomical and each volume of the functional data set to the lowest motion volume. We also blurred each volume to 4mm fwhm and censored our data, removing runs where participants moved more than 3mm. Multiple linear regression was then applied, allowing for contrasts between moving and static blocks. hMT+ and other motion processing regions were localized using a moving > static dots contrast in the contralateral visual field.

### 6. Stimulation parameters and positioning

We delivered stimulation using a battery-driven stimulator (NeuroConn DC STIMULATOR MC) and a pair of saline-soaked sponge electrodes (NaCl concentration: 0.9%). We stimulated using hf-tRNS at 1.5 mA (crest to trough) alternating current with 0-offset and random frequencies at the capacity of our stimulator (range between 101 to 640 Hz). For the sham condition, we applied the electrodes over hMT+ and took impedance exactly as done for active stimulation, but then did not turn the stimulation on^1^. Stimulation lasted for a total of 18 minutes. Each electrode had an area of 25 cm^2^. We kept the current density below safety limits (below 1 A/m^2^).

In the experimental and sham conditions, we placed one electrode over the left hMT+, and in the active control condition, over the forehead. The other electrode in all conditions was placed over the vertex (Cz). Left hMT+ was localized using predetermined coordinates, 3 cm dorsal of the inion and 5 cm leftward (Ghin et al., 2018). The vertex was identified using Cz from the International 10-20 system, as in Ghin et al. (2018). Stimulation differences to the original Ghin et al., (2018) study are listed in Table 1.

For the control stimulation, we placed one electrode over the left forehead, as described by Ghin et al. (2018). Additionally, using Brainsight to provide MNI coordinates for each participant’s forehead stimulation location, we simulated the stimulation e-fields with ROAST (Huang et al. 2019). Using these simulations, the MNI coordinate with the largest e-field value was located in the frontal lobe (x=-27.61(SD=5.32), y=56.15(SD=4.02), z=1.22(SD=5.98).

Examining a random 25 articles of the 203 entries from Neurosynth.org (NeuroSynth, RRID:SCR_006798) which report these MNI coordinates, the regions most impacted were dorsolateral prefrontal cortex (DLPFC), middle frontal gyrus, and orbital frontal cortex. These articles reported that activity in these regions is related to working memory tasks, reward learning and semantic tasks. Table S1 list all 25 randomly-selected articles, the associated brain regions, and tasks performed in the scanner.

Participants were blinded to the type of stimulation applied in each session. To assess blinding integrity, we asked participants if they perceived any sensation (tingling, burning or pain) under the electrodes after each session. Sensation experience provides a measure for the perception of present or absent stimulation. Participants reported a perceived sensation during stimulation 9.5% of the time for the active hMT+ condition, 11.9% of the time for the sham hMT+ condition, and 9.5% of the time for the forehead control condition. Additionally, in the final session we also asked participants to report if they thought they received stimulation in that session and how confident they were in their report. This report enabled us to determine that participants cannot distinguish between sham, active stimulation targeted at hMT+, and active stimulation to the forehead in a between-subjects comparison. These questions evoke a response about the perception of stimulation, reserving the direct question about stimulation presence to the end of the experiment and avoiding preconceptions about the presence of stimulation during the experimental protocol. Of the 14 participants whose final session was active hMT+ stimulation, 46% reported believing they received stimulation (six yes, seven no, one data not collected); of the 14 participants whose final session was sham over hMT+, 46% reported believing they had received stimulation (six yes, seven no, one data not collected); and of the 14 participants whose final session was stimulation delivered to the forehead, 36% reported believing they had received stimulation (five yes, nine no).

### 7. Procedure

Our within-subjects design required each participant to receive hf-tRNS targeted to the hMT+ and forehead, and sham stimulation to the hMT+ for a total of three non-consecutive sessions. There was a minimum of 72 hours between each stimulation session for each participant. We counterbalanced condition order to account for learning effects. Since we are interested in the effects of online stimulation, participants performed the task during stimulation. Each participant underwent a stimulation session consisting of 160 trials in each visual field split across five blocks. MLPs were run on trials presented in each visual hemi-field (left and right).

Over the course of one stimulation session, the coherence threshold for the left and right hemi-fields was estimated from MLPs across all 160 trials. In addition to the stimulation sessions, participants were brought in for a fMRI session to localize regions active for lateralized motion processing, including left hMT+ (Figure 3). These data were used for our exploratory analysis comparing the activity maps of lateralized motion processing to anatomically defined stimulation sites.

**Figure 3.**
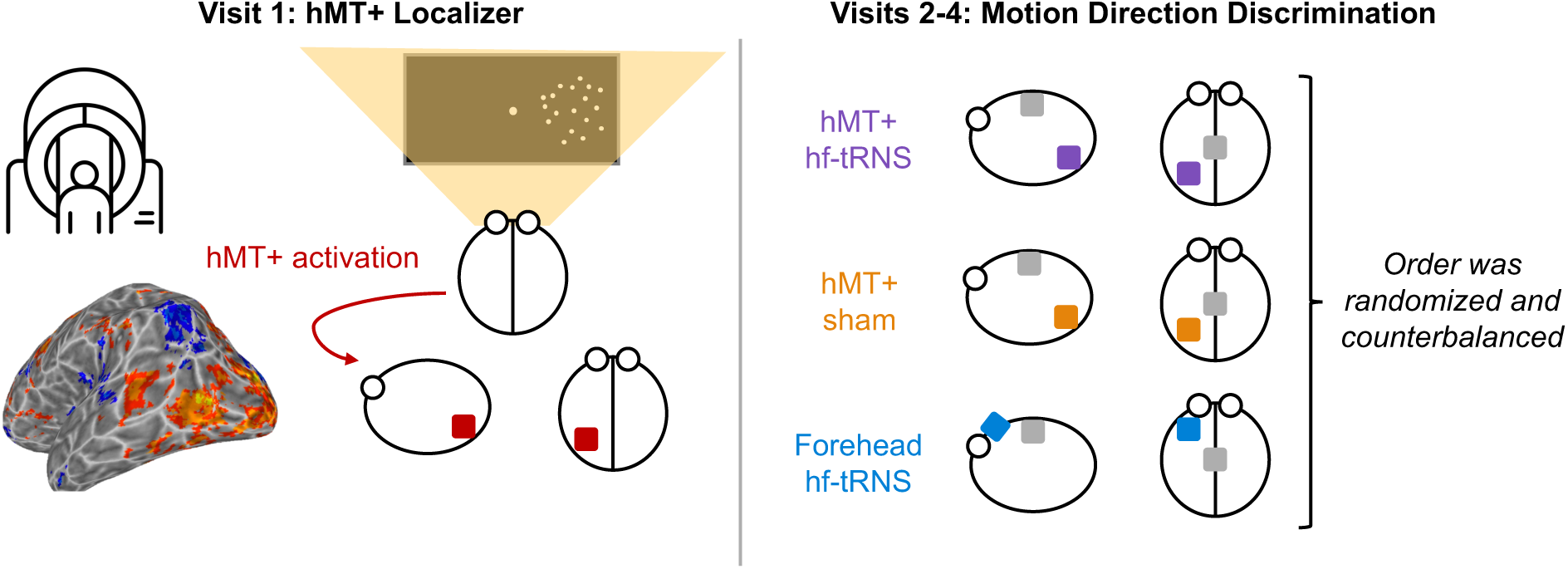
Experimental procedure according to visit. Visit 1 consisted of a fMRI hMT+ localizer (bottom left: localized left hMT+ (right motion > static)). For Visits 2-4, participants performed the motion discrimination task under three different conditions. These were the experimental condition, hMT+ stimulated with hf-tRNS, and two control conditions: i) hMT+ with sham stimulation, and ii) forehead stimulated by hf-tRNS. The order of Visits 2-4 was randomized and counterbalanced.

We hypothesized that in the right visual field (contralateral to stimulation) the coherence threshold would be facilitated by hf-tRNS stimulation to the left hMT+, as was found by Ghin et al. (2018). We further hypothesized that neither stimulation to the forehead nor sham stimulation would modulate the coherence threshold.

### 8. Statistical analysis

We performed three one-tailed paired sample *t*-tests to examine:

1. the decrease in motion coherence threshold to the visual field contralateral from stimulation in comparison to the visual field ipsilateral from stimulation following hf-tRNS to hMT+.
2. the difference between contralateral versus ipsilateral motion coherence thresholds following hMT+ targeted hf-tRNS versus the difference between contralateral versus ipsilateral motion coherence thresholds following sham hf-tRNS.
3. the difference between contralateral versus ipsilateral motion coherence thresholds following hMT+ targeted hf-tRNS versus the difference between contralateral versus ipsilateral motion coherence thresholds following forehead targeted hf-tRNS.

### 9. Hypothesized results

We hypothesized we would find a reduction in coherence threshold for hf-tRNS stimulation to the left hMT+ in the contralateral right visual field in comparison to ipsilateral left visual field. We also hypothesized the facilitation of contralateral in comparison to ipsilateral motion coherence threshold to be larger for hMT+ targeted hf-tRNS in comparison to hMT+ sham tRNS and for hMT+ targeted hf-tRNS in comparison to forehead targeted hf-tRNS.

### 10. Exploratory overlap analyses

In an exploratory analysis, we compared the fMRI data with the simulated e-field for the anatomically targeted hMT+ in native brain space for each participant. Our goal was to determine if the functionally localized right visual field motion processing regions overlapped closely with the strongest e-field values. Specifically, we predicted a greater extent of overlap would predict a larger impact of hf-tRNS on motion processing in the right visual field. To prepare the functional MRI data, we used the right moving dot > right static dot contrast and extracted the t-statistic across the whole brain (See *fMRI analysis and localization of the hMT+)*. To run the e-field simulation of the anatomically defined hMT+, we localized the MNI coordinates of the stimulation site using Brainsight (v2.4.11) at the time of stimulation and ran the ROAST toolbox to simulate the e-field for the active left hMT+ montage (between anatomically localized hMT+ and Cz) for each individual. Finally, we ranked the voxels from highest to lowest t-statistic and highest to lowest e-field value separately and then quantified the mutual information between the two datasets (Giangregorio, 2022). No threshold was applied to the fMRI data.

### 11. Data & Code Availability

All data is available on OSF (https://osf.io/chx6z/). To access the data, navigate to the Files tab, and Data. The code for running the experimental paradigm and motion localizer, and analyzing the data are available on GitHub (https://github.com/gcaedwards/tRNS_hMT_RegisteredReport).

## Results

### Hf-tRNS to left hMT+ does not improve motion processing

Figure 4A shows the group mean motion coherence thresholds, separated by hemifield, for each of the three stimulation conditions. To answer our first hypothesis, we first performed a 1-tailed paired samples *t*-test to determine if the ipsilateral (mean=70.69 and SEM=3.15) motion coherence threshold was larger than the contralateral motion coherence threshold (mean=71.86 and SEM=3.40), revealing no significant difference (t(41)=-0.61,p=0.72; one-tailed). Therefore, we found no evidence that hf-tRNS to left hMT+ improved motion coherence processing in the contralateral right visual field only.

**Figure 4.**
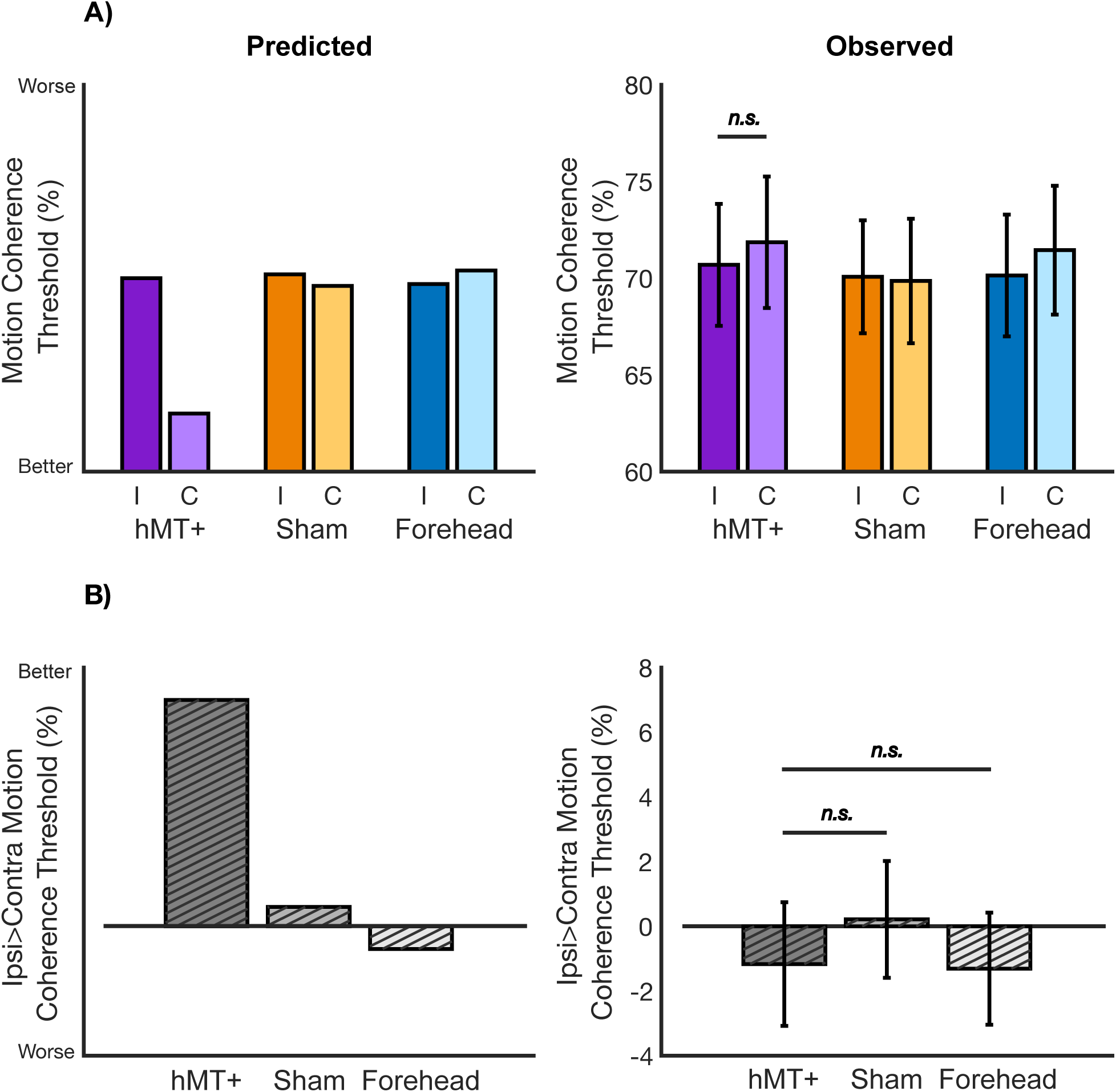
Predicted and observed results. 4A) Mean coherence thresholds for each stimulation condition, split by the ipsilateral and contralateral visual fields. Left: we predicted a significant drop in motion coherence threshold for hMT+ stimulation in the contralateral visual field only. Right: we demonstrated no decrease in motion coherence threshold in the contralateral visual field for hMT+ stimulation. 4B) Mean ipsilateral versus contralateral motion coherence threshold difference. Left: we predicted the difference between ipsilateral and contralateral motion coherence threshold to be significantly higher in the active hMT+ condition compared with sham and forehead control. Right: we demonstrated active hMT+ stimulation was no different from sham or forehead control in modulating contralateral versus ipsilateral motion coherence threshold. Error bars are standard error across participants, and n.s. denotes non-significant comparison.

Approaching hypotheses two and three, we calculated the ipsilateral versus contralateral motion coherence difference (hMT+ mean=-1.17 and SEM=1.91; Sham mean=0.21 and SEM=1.81; Forehead mean=-1.31 and SEM=1.73; Figure 4B). We found no difference between active hMT+ stimulation and sham hMT+ (t(41)=-0.72,p=0.76), and no difference between active hMT+ stimulation and active Forehead stimulation (t(41)=0.06,p=0.47). Therefore, active hMT+ stimulation did not modulate motion coherence processing in the contralateral field any differently to our two within-subject control conditions.

### Exploratory Results

#### No visual field-wide stimulation effect

We used larger electrodes over hMT+ in comparison to Ghin et al., 2018, prompting the consideration of a whole field motion effect rather than contralateral only. Specifically, our larger electrodes may have been less targeted and significantly modulated MST, a motion processing region with receptive fields which extend into the ipsilateral as well as contralateral visual fields. We averaged performance for the ipsilateral and contralateral visual field of hf-tRNS to hMT+, sham, and the forehead and fit a repeated measures model. We found no significant difference between these conditions after collapsing across visual field (F(2)0.18, p=0.83).

#### Extent of mutual information between functionally localized motion processing regions and simulated active hMT+ e-field does not predict behavior

We ranked the t-statistic fMRI voxels and e-field simulation voxels from highest to lowest separately and then quantified the mutual information between the two datasets (Giangregorio, 2022). The mutual information calculated between the functionally localized right visual field motion processing activity and simulated active hMT+ e-field (Figure 5A & 5B, respectively) is presented as a percentage of maximum possible overlap, which differs for each participant (due to differing brain characteristics). Note, one participant was excluded from the analysis due to corruption of simulation results. We found no correlation between mutual information values and the ipsilateral>contralateral behavioral difference across participants (Pearsons correlation = −0.24, p = 0.13); Figure 5C). Therefore, the strength of the overlap between the functionally localized rightward motion processing brain regions and simulated e-field for hMT+ stimulation did not predict a stronger influence of the stimulation on behavior. This analysis undermines the suggestion that functionally localized stimulation would have improved the stimulation impact on contralateral motion processing. To support this analysis, we also correlated the voxel ranking of the functional localizer and e-field simulation which also did not predict behavior following hMT+ stimulation (Supplemental Figure 1; Pearsons correlation = −0.01, p = 0.95). Further, we examined the MNI distance (mm) between the anatomically localized and functionally localized hMT+ and found no relationship with behavioral modulation from active hf-tRNS to hMT+ (Supplemental Figure 1; Pearsons correlation = −0.02, p = 0.90). Note, two participants were excluded from the MNI distance analysis due to poor localization of hMT+.

**Figure 5.**
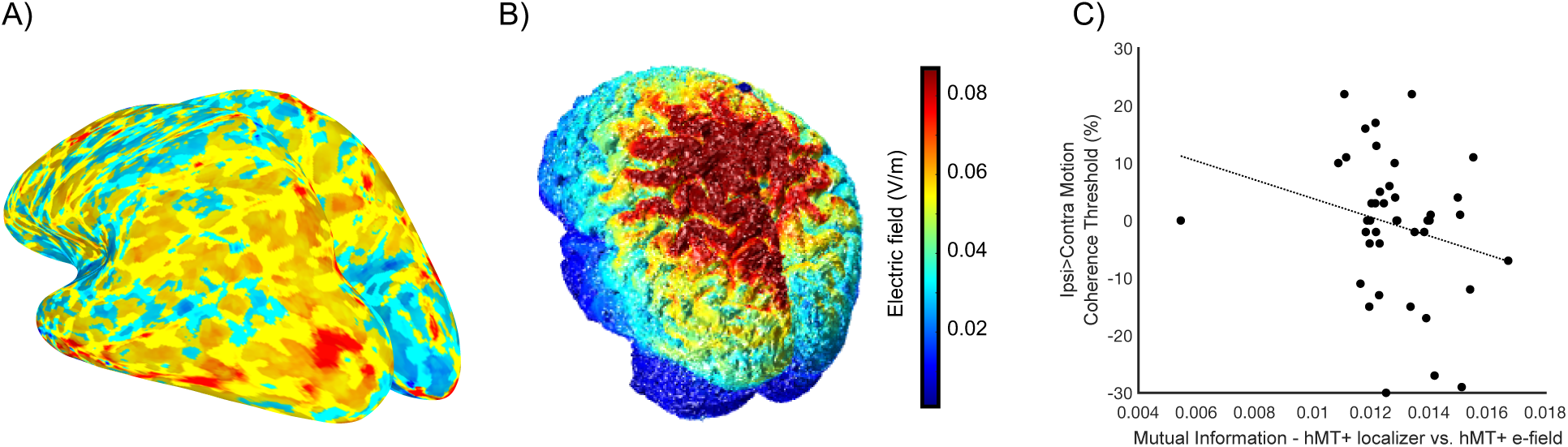
Mutual information between the functionally localized motion processing regions and simulated active hMT+ to Cz stimulation. 5A) Example participant functional localizer using contrast moving > static dots contrast in the contralateral visual field. No threshold applied. 5B) Same example participant e-field simulation for hMT+ to Cz. 5C) Mutual information of functionally localized left hMT+ (5A) and e-field simulation (5B) correlated with ipsi>contralateral motion coherence threshold following hMT+ active stimulation. Least-squares line plotted.

## Discussion

We found no impact of hf-tRNS targeted at left hMT+ on a contralateral motion discrimination task, failing to reproduce an effect from a previous study (Ghin et al., 2018). Specifically, in comparison to the ipsilateral visual field, motion processing in the contralateral visual field from left hMT+ stimulation was not modulated. Further, there was no difference between active hMT+ stimulation and sham over hMT+, or the active control over left forehead. In attempting to reproduce the effects from Ghin et al. (2018), we made minor adjustments to the protocol to target our replication of interest. Table 1 lists the methodological differences between the original Ghin et al., (2018) study and the present study. First, we combined stimulation conditions from Ghin et al.’s experiment 1 (hf-tRNS to hMT+, sham stimulation to hMT+) and control experiment 2 (hf-tRNS to forehead control). Examining all conditions within-subject should have increased sensitivity to neuromodulation through accounting for inter-subject variability (Greenwald, 1976), however we were unable to detect our predicted behavioral modulation. While it is possible that including more within-subject conditions could have resulted in significant learning effects across session, which increases noise and reduces the chance to detect stimulation effects, our final number of within-subject behavioral sessions was four (including the thresholding), the exact same number of sessions in Ghin et al., Experiment 1. Specifically, Ghin et al., Experiment 1 originally included two tDCS sessions as well as the tRNS and sham sessions. We did not include the tDCS due to the lack of modulation detected (Ghin et al., 2018). We also deviated from the original protocol by including more participants indicated by power analyses. More participants should also reduce the impact of participant variability on the group level analyses, enabling a more reliable result. Our sample size was selected to achieve a 0.9 power to test our minimum effect size of interest based on the data from Ghin et al., (2018). We predicted larger between-subject variance in Ghin et al. and therefore felt confident this was a conservative estimate for our within-subject design. However, it is plausible our sample would not have been large enough to detect a potentially smaller effect size.

One protocol adjustment which could have contributed to differential outcomes between our current study and the original Ghin et al. (2018) findings was our use of differently sized electrodes. Ghin et al., (2018) targeted hMT+ and the forehead with 16cm^2^ sized electrodes, with the second electrode over Cz sized 60cm^2^, while we used 25 cm^2^ electrodes for both electrode sites in each stimulation visit. The different electrode sizes may have contributed to differences in the electric field applied to the cortex. hMT+ the target region of our stimulation, contains a strong representation of motion in the contralateral visual field (Strong et al., 2017; Ajina et al., 2015; Braddick et al., 2001). Within hMT+ lies MT, which has a more dominant contralateral visual field motion representation, and MST, a motion sensitive visual processing region which contains receptive fields that extend into the ipsilateral visual field (Amano et al., 2009). It is possible that our larger electrodes at the target site captured MST, while Ghin et al. were able to selectively target MT, enabling the stimulation effect difference between visual fields that we do not observe in our data. However, we would have expected to observe a general improvement in performance across both visual hemifields in our active condition compared to sham in the case that MST was also stimulated. A visual field wide improvement between hf-tRNS to hMT+ and sham to hMT+ is not observed in our data.

The maximum likelihood procedure (MLP) thresholding over all five blocks could also have impacted the result. Resetting the threshold for each block like Ghin and colleagues did, could lead to a different result than our approach of the thresholding performed across all five blocks. We chose to threshold across all five blocks to allow the MLP to operate on as much data as possible. Working across more trials, the MLP should have generated a highly reliable estimate of the motion coherence level at which each participant performs the task at 70% accuracy. Thus, the different thresholding procedures could yield a difference in threshold estimate. However, we would still expect to observe a quantitative difference between the ipsilateral and contralateral hemifields, which we do not.

Critically, if the minor protocol adjustments outlined above impact hf-tRNS over hMT+, it brings into question the efficacy of hf-tRNS in motion discrimination. For hf-tRNS to be beneficial in clinical and research situations, a replicable effect which is robust to small changes and advances in procedures are important.

One major direction the non-invasive brain stimulation field is taking is individualized stimulation protocols. An approach for individualized stimulation is the localization of each individual’s stimulation site based on functional neuroimaging rather than averaged anatomical coordinates. With the goal of replicating Ghin et al., (2018) as closely as possible, we used the anatomical landmark of the inion and measured to our stimulation target using 3 cm dorsal of the inion and 5 cm leftward (typical of stimulation targeted at hMT+ without neuroimaging support; Ghin et al., 2018; Campana et al., 2002; 2006). In addition, we collected a functional localizer to determine if the simulation of the e-field from the anatomically localized hMT+ was well aligned to the functional data for localized hMT+. We found that even with a close alignment, we could not predict an improved behavioral outcome. This exploratory analysis enables some certainty that functionally localized hMT+ would not have changed the stimulation outcome.

Even though we were unable to replicate the findings of Ghin and colleagues, there is evidence from other empirical work that hf-tRNS has a beneficial effect on behavior. Many protocols with a successful facilitatory impact on behavior are run with a tRNS intensity of 1 mA peak-to-trough (Romanska et al., 2015; Contò et al., 2021; van der Groen & Wenderoth, 2016). In fact, one previous replication failure included a change in stimulation intensity from success at 1 mA (Romanska et al., 2015) to lack of replication using 1.5 mA (Willis et al., 2019). van der Groen & Wenderoth (2016) demonstrated that the optimal stimulation range for hf-tRNS is 0.75-1 mA peak-to-trough, suggesting our stimulation intensity may have been out of range for facilitatory effects. The stochastic resonance theory of tRNS suggests random noise added to the brain enhances a subthreshold neural signal, resulting in an improvement in behavior. 1.5 mA may introduce too much noise into the system, masking neural firing patterns, culminating in a negative impact on behavioral performance (van der Groen & Wenderoth, 2016).

Additionally, it may be the case that bilateral tRNS protocols, which involve delivering stimulation to homologous regions on either side of the brain, are more efficacious. Therefore, the present design targeting hMT+ unilaterally may be insufficient to demonstrate a robust behavior modification resulting from stimulation. However, this is an empirical question and requires new studies to examine the impact of bilateral versus unilateral hMT+ stimulation in motion discrimination tasks.

Finally, tRNS has been recommended for clinical application due to the previous evidence of strong behavioral effects, and participants inability to sense the stimulation. Tolerance of interventions for clinical populations is critical for intervention success. Here, we confirm that our participants were unable to determine if they received active stimulation (over hMT+ or the forehead) or sham.

Ultimately, we were unable to replicate motion discrimination facilitation following hf-tRNS targeted at hMT+. Our failure to replicate the original finding undercuts the generalizability of tRNS as a technique. Moving forward, more research will be necessary to demonstrate which montages and protocols produce successful replications of tRNS to show translational value for modulating behavior.

## Supplemental

**Supplemental Figure 1.**
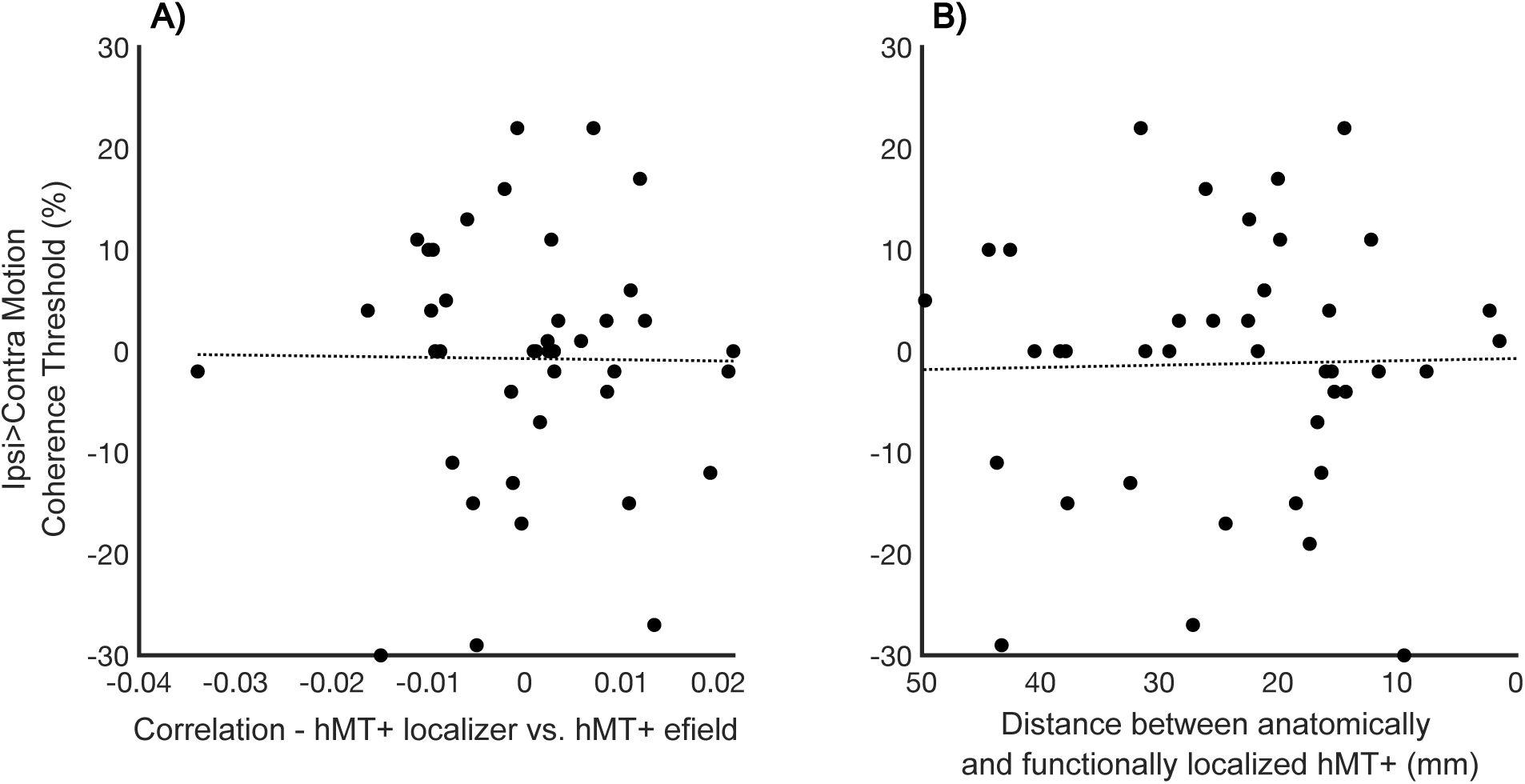
S1A) Correlation of functionally localized left hMT+ and e-field simulation used to predict ipsi>contralateral motion coherence threshold following hMT+ active stimulation. S1B) Distance between functionally localized left hMT+ and anatomically localized hMT+ correlated with ipsi>contralateral motion coherence threshold following hMT+ active stimulation. Least-squares line plotted.

**Supplemental Table 1.** Lists 25 of 203 articles associated with the MNI coordinates with the largest e-field value located in the frontal lobe (x=-27.61, y=56.15, z=1.22) from Neurosynth.org (NeuroSynth, RRID:SCR_006798)

| Article | Region | Task |
| --- | --- | --- |
| Alvarez, R. P., et al., (2011) | Middle/superior frontal gyrus | Fear conditioning |
| Bereczkei, T., et al. (2013) | Middle/superior frontal gyrus | Trust game |
| Cheke, L. G., et al., (2017) | Dorsolateral prefrontal gyrus (DLPFC) | What-where-when episodic memory task |
| Collins, A. G. E., et al., (2017) | Middle frontal gyrus | Reward learning |
| Dove, A., et al. (2006) | Left superior frontal gyrus | Memory encoding/retrieval |
| FeldmanHall, O. et al. (2015) | Left orbital frontal cortex | Pain vs. Gain task |
| Forbes et al., (2012) | Orbital frontal cortex | Spatial task |
| Ford et al., (2011) | Frontopolar prefrontal cortex | Autobiographical memory retrieval |
| Greene et al., (2004) | Middle Frontal Gyrus, DLPFC | Personal moral judgement |
| Hartogsveld et al., (2018) | Lateral frontal pole | Relational processing |
| **Hermans et al., (2008)** | Orbitofrontal cortex, BA47 | Facial expression viewing |
| **Jenkins et al., (2014)** | Left middle frontal gyrus | Theory of mind task |
| **Kalpouzos et al., (2009)** | Left Frontal mid/mid orb/sup (10, 47) | Episodic memory encoding |
| **King et al., (2005)** | Anterior prefrontal cortex | Contextual episodic memory task |
| **Limb & Braun (2008)** | Polar medial prefrontal - middle | Piano improvisation |
| **Longe et al., (2009)** | Lateral PFC | Memory load |
| **Manelis et al., (2017)** | Frontal Pole | Temporal memory |
| **Olsson et al., (2007)** | Middle frontal gyrus | Social fear transmission |
| **Pyke et al., (2017)** | Brodmann Area 10 | Typing in a keypad when either visuospatially trained or symbolically trained |
| **Schon et al., (2008)** | Left Frontomarginal sulcus (anterior VLPFC) | Delayed match to object or place |
| **Specht et al., (2009)** | Middle frontal gyrus | WCST |
| **Wendelken et al., (2008)** | Rostrolateral PFC | Delayed item recognition |
| **Winston et al., (2014)** | Left frontomarginal sulcus | Pain valuation task |
| **Wu et al., (2009)** | Middle frontal gyrus | Semantic task |
| **Zanto et al., (2016)** | Middle frontal orbital | Memory task with face stimuli |

## Appendix A: Simulations of stimulation montages

**Appendix A, Figure 1:**
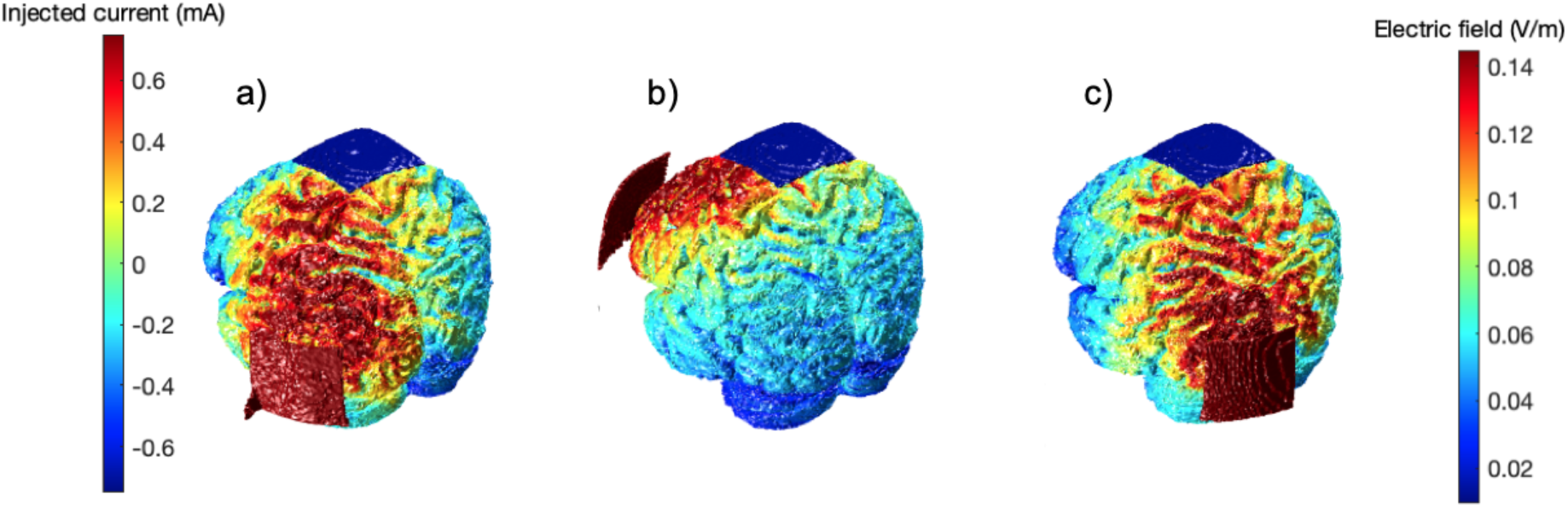
Simulations of a) T5 (left hMT+) to Cz, b) Fp1 (left forehead) to Cz, c) O1 (left V1) to Cz using ROAST (Huang et al., 2019).

The ROAST toolbox was designed for quasi-static modelling when the frequency is lower than 1k Hz. The head model is ohmic for the purpose of the simulation and is therefore the same for direct or alternating current (Huang et al., 2019).

Matlab Syntax:

**Left hMT+ to Cz:** roast(’example/subject1.nii’, {’T5’,0.75,’Cz’,-0.75}, ‘electype’, {’pad’,’pad’}, ‘elecsize’, {[50,50,3],[50,50,3]}, ‘simulationTag’, ‘hMTsimulation’)

**Left forehead to Cz:** roast(’example/subject1.nii’,{’Fp1’,0.75,’Cz’,-0.75}, ‘electype’,{’pad’,’pad’}, ‘elecsize’,{[50,50,3],[50,50,3]}, ‘simulationTag’, ‘foreheadsimulation’)

**Left V1 to Cz:** roast(’example/subject1.nii’,{’O1’,0.75,’Cz’,-0.75}, ‘electype’,{’pad’,’pad’}, ‘elecsize’,{[50,50,3],[50,50,3]}, ‘simulationTag’, ‘V1simulation’)

The active control of V1 to Cz as used by Ghin et al. (2018) was excluded from replication due to the overlap between the simulated electric field with the simulation of hMT+ to Cz. Overlap was quantified by analyzing the mutual information in the hMT+ and V1 simulations in comparison to the mutual information in the hMT+ and forehead simulations (Giangregorio, 2022). Where zero would denote complete independence between the simulations, and 2.50 complete dependence, hMT+ and V1 had a mutual information score of 1.30, whereas hMT+ and the forehead had a mutual information score of 0.95. Whilst some dependence is expected due to the same reference electrode position (Cz) in all montages, minimizing mutual information was preferable.

## Appendix B: Power analyses

We performed three power analyses on our three comparisons of interest using a plot digitizer to extract the data from the figures:

First, we examined contralateral versus ipsilateral motion coherence thresholds following hf-tRNS to hMT+. Specifically, we used a mean of 28.75% related to the contralateral motion coherence threshold, a mean of 39.26% related to the ipsilateral motion coherence threshold, and standard deviation of 17.56% related to the standard deviation in the ipsilateral visual field following hf-tRNS targeted at left hMT+ (Ghin et al., 2018 Figure 2a; Cohen’s d: 0.60). With a power of 0.9 and type I error rate of 2%, we calculated the need for 34 participants.

Second, we examined the difference between contralateral versus ipsilateral motion coherence thresholds following hMT+ targeted hf-tRNS versus the difference between contralateral versus ipsilateral motion coherence thresholds following sham hf-tRNS. We used a mean of 10.51% related to the difference in contralateral and ipsilateral motion coherence threshold for hMT+ targeted hf-tRNS, a mean of 2.59% related to the difference in contralateral and ipsilateral motion coherence threshold for hMT+ targeted sham tRNS, and standard deviation of 14.8% related to the standard deviation in the contralateral visual field following sham tRNS targeted at left hMT+ (Ghin et al., 2018 Figure 2a; Cohen’s d: 0.54). With a power of 0.9 and type I error rate of 2%, we calculated the need for 42 participants. Finally, we examined the difference between contralateral versus ipsilateral motion coherence thresholds following hMT+ hf-tRNS versus the difference between contralateral versus ipsilateral motion coherence thresholds following forehead targeted hf-tRNS. We used a mean of 10.51% related to the difference in contralateral and ipsilateral motion coherence threshold for hMT+ targeted hf-tRNS, a mean of 1.17% related to the difference in contralateral and ipsilateral motion coherence threshold for forehead targeted hf-tRNS, and standard deviation of 8.44% related to the standard deviation in the contralateral visual field following hf-tRNS targeted at the left forehead (Ghin et al., 2018 Figures 2a and 4a; Cohen’s d: 1.11). With a power of 0.9 and type I error rate of 2%, we calculated the need for 12 participants. To maximally power our study for our three effect sizes of interest, we collected 42 participants.

## Appendix C: Screening participants for 70% motion discrimination

During piloting (see below), we were unable to establish a threshold at 70% correct on some participants due to poor performance. A threshold target of 70% is necessary for the maximum likelihood procedure thresholding to function. In order to replicate the original findings, rather than reduce the p-target, we screened participants using a non-adaptive constant thresholding procedure.

### Piloting the RDKs with non-adaptive constant thresholding

Using the same random dot kinematograms (RDKs) presented in motion discrimination task as described in the Stimuli section of the Method, we presented eight blocks of RDKs (n trials = 64) at a range of motion coherence levels from 25% to 100% in steps of 5%. This resulted in 32 trials per motion coherence level across all blocks. Using these data, we fit a psychometric function using a logit glm, and find the motion coherence at which participants were performing at 70% correct.

We plot these functions for nine pilot participants in Appendix C, Figure 1. Five performed the motion discrimination task above 70% at one or more of the motion coherence levels, the other four performed below 70% at all motion coherence levels.

**Appendix C, Figure 1:**
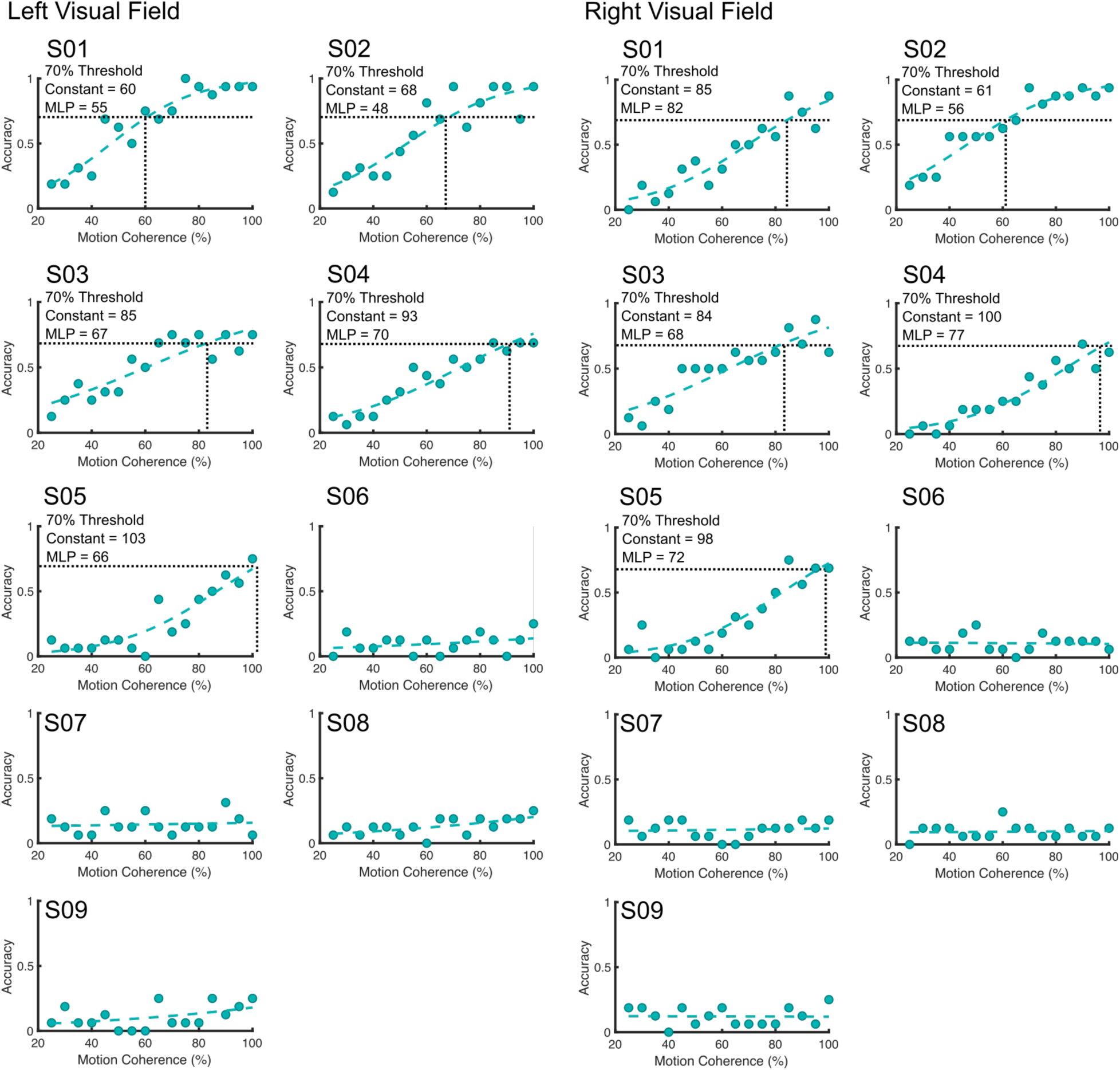
Pilot data of nine participants in left and right visual fields. Data from the constant thresholding plotted with a psychometric function and MLP reported.

The five pilot participants able to perform the task above 70% correct returned for a second session using the MLP, and we found the MLP provided estimated thresholds lower than the constant thresholding performed in the first session, which may reflect a session effect of learning or increased sensitivity of the MLP approach.

### Screened participant data

In Appendix C, Figure 2 we present all screened participants (1-42 included in the study; 43-87 excluded). Participants were excluded due to subthreshold performance at all motion coherence levels.

**Appendix C, Figure 2:**
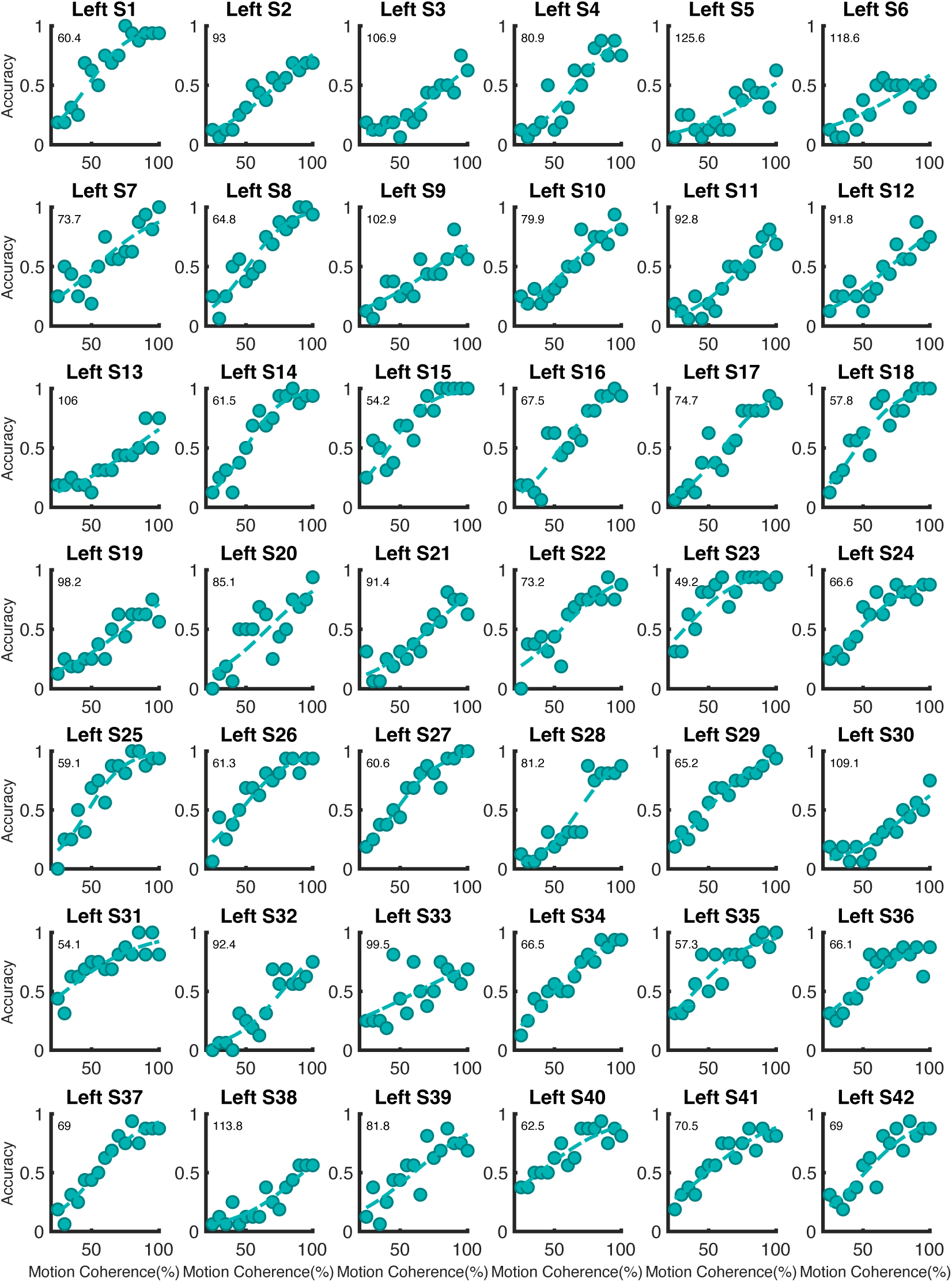
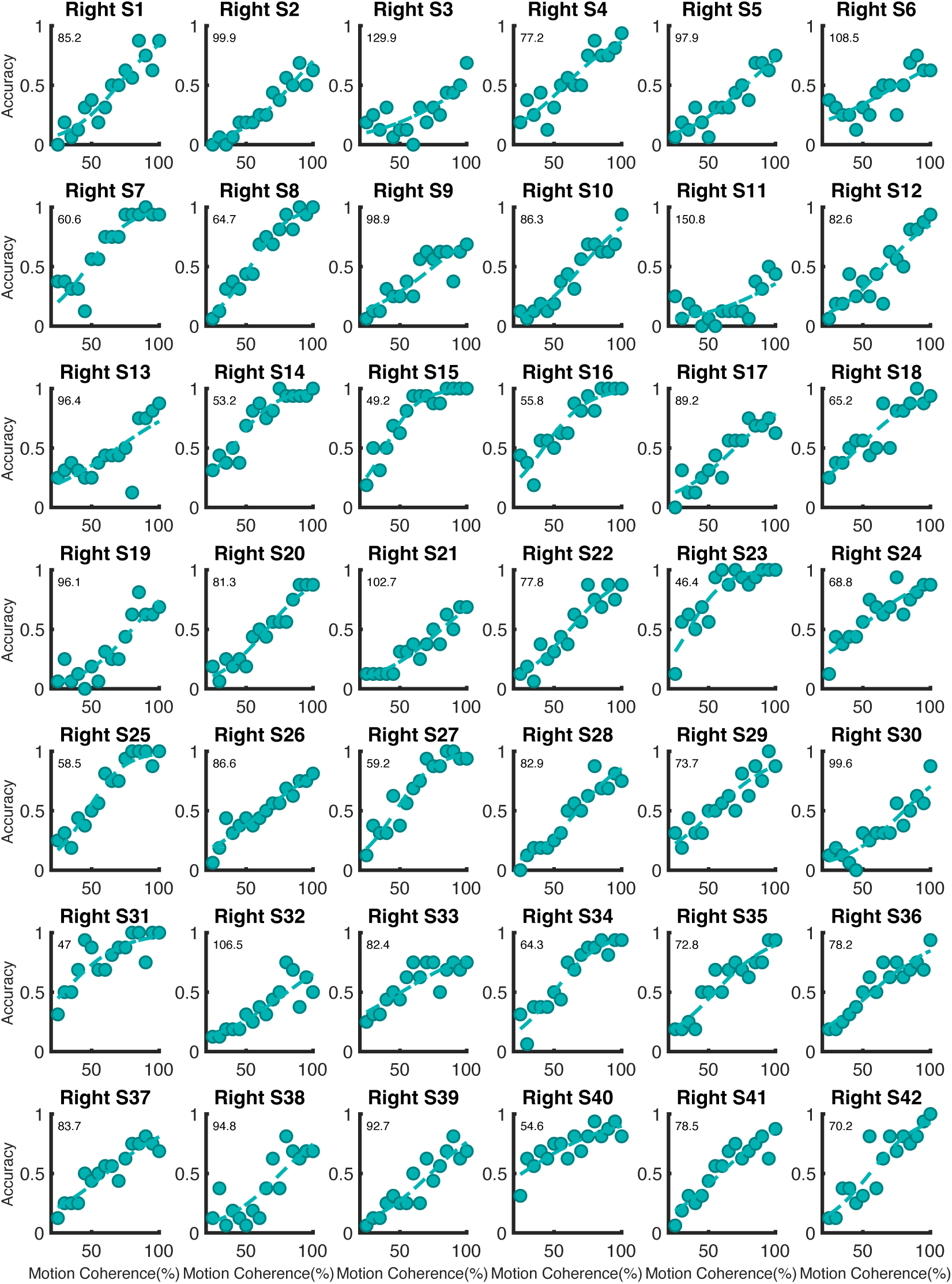
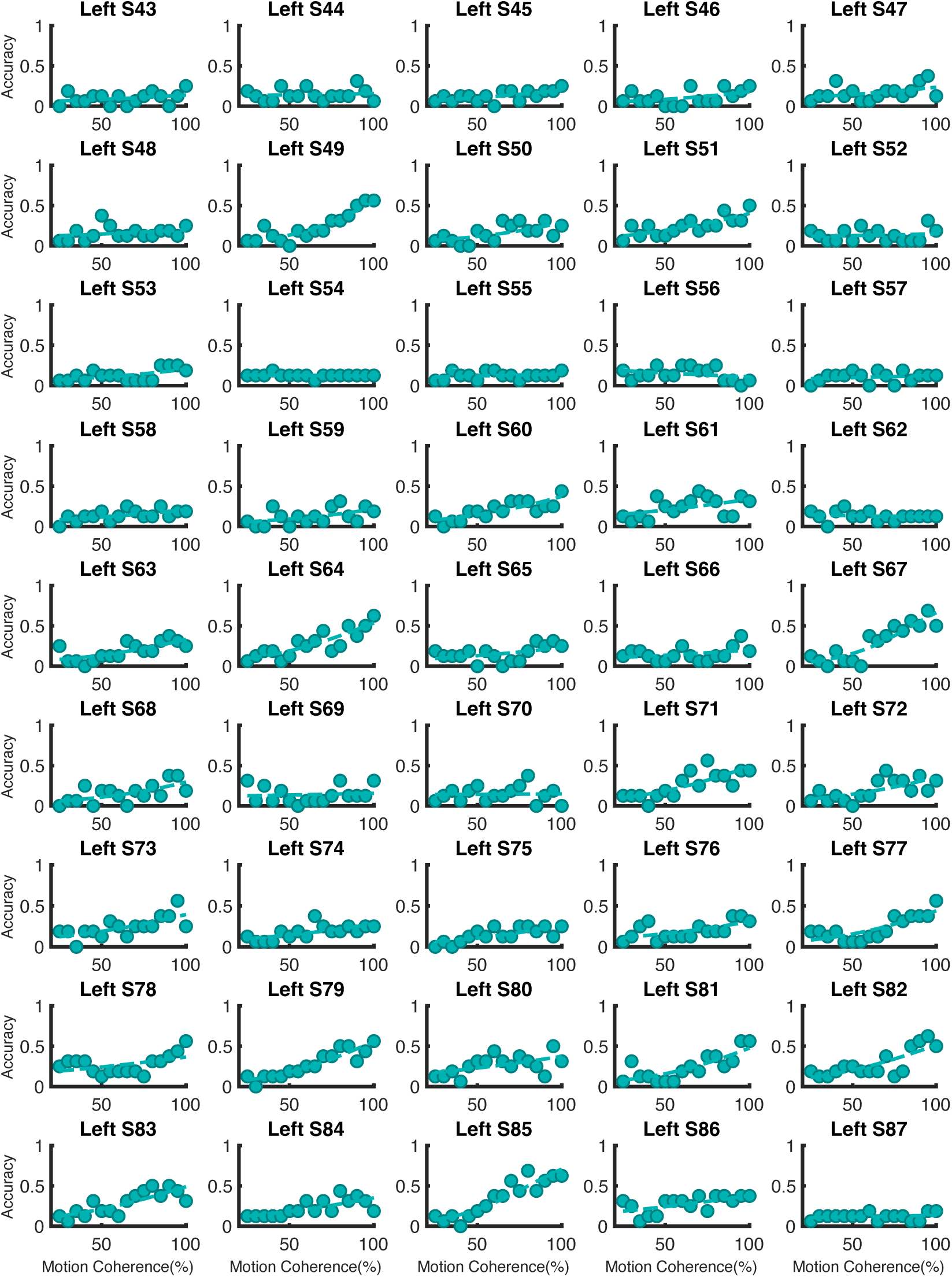
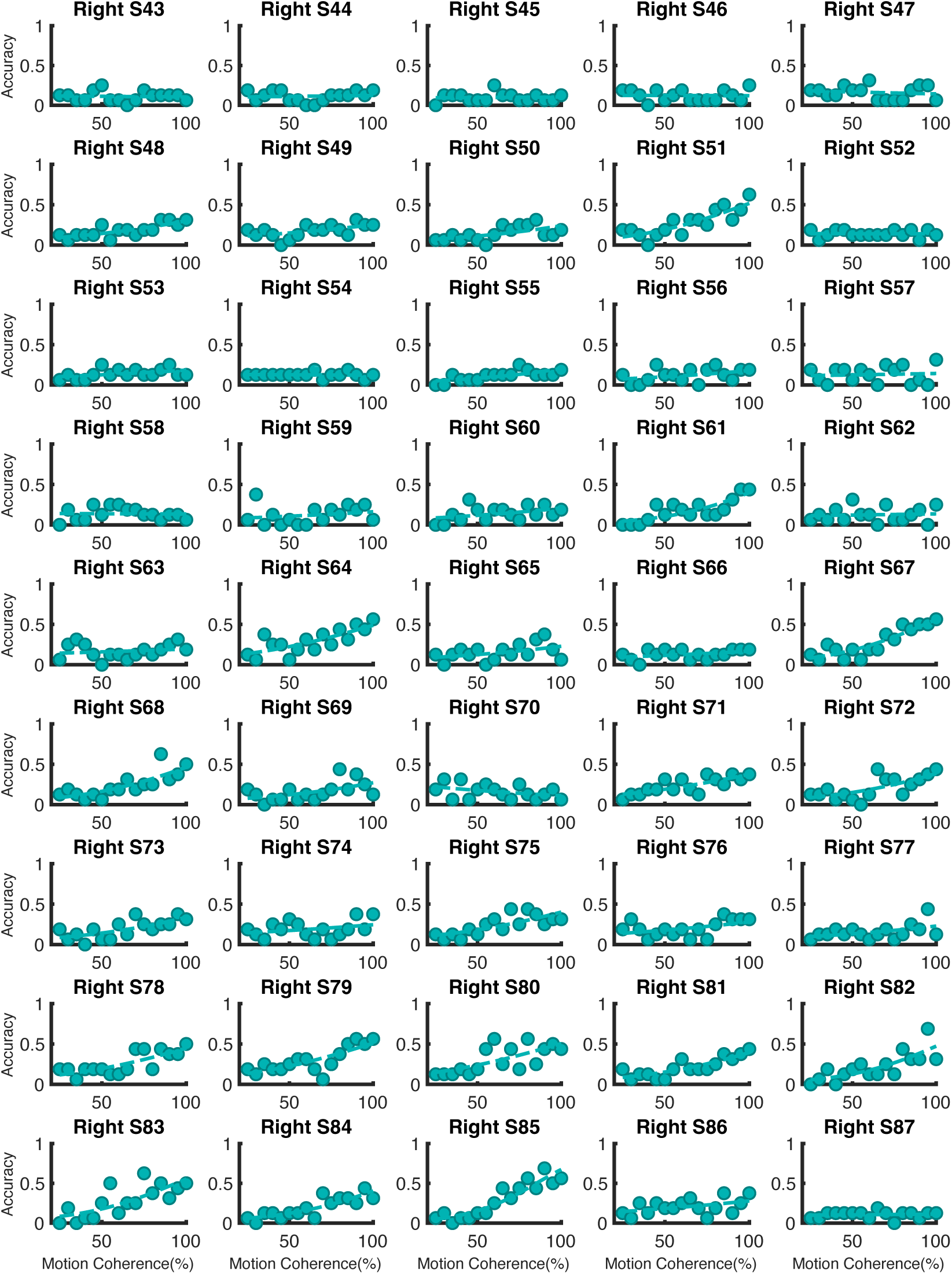
Screening data of 87 participants in left and right visual fields. Data from the constant thresholding plotted with a psychometric function and constant threshold reported for participants 1-42 who were included in the main study.

## Acknowledgements

This research was supported by the Intramural Research Program of the NIMH (ZIAMH002909 & ZIAMH002893). The ClinicalTrials.gov identifiers are NCT01617408 & NCT00001360.

## Footnotes

1 In the In Principle Stage 1 submission we registered that the sham condition would be ramped up for 30 seconds, but due to user error no such ramp up was performed.

## Notes

https://osf.io/chx6z/

https://github.com/gcaedwards/tRNS_hMT_RegisteredReport

### Competing Interest Statement

The authors have declared no competing interest.

### Summary of Updates

Badge added to denote a positive recommendation from Peer Community In Registered Reports (PCI RR)

